# Zn^2+^ intoxication of *Mycobacterium marinum* during *Dictyostelium discoideum* infection is counteracted by induction of the pathogen Zn^2+^ exporter CtpC

**DOI:** 10.1101/575217

**Authors:** Louise H. Lefrançois, Vera Kalinina, Elena Cardenal-Muñoz, Nabil Hanna, Hendrik Koliwer-Brandl, Joddy Appiah, Florence Leuba, Hubert Hilbi, Thierry Soldati, Caroline Barisch

## Abstract

Macrophages use diverse strategies to kill or restrict intracellular pathogens. Some of these strategies involve the deprivation of bacteria from (micro)nutrients such as transition metals, and the bacteria intoxication through metal accumulation. Little is known about the chemical warfare between *Mycobacterium marinum*, a close relative of the human pathogen *M. tuberculosis*, and its hosts. Here we use the professional phagocyte *Dictyostelium discoideum* to investigate the role of Zn^2+^ during *M. marinum* infection. We show that *M. marinum* infection induces the accumulation of Zn^2+^ inside the *Mycobacterium*-containing vacuole (MCV), achieved by the induction and recruitment of the *D. discoideum* Zn^2+^ efflux pumps ZntA and ZntB. In cells lacking the ZntA detoxifying transporter there is further attenuation of *M. marinum* growth, possibly due to a compensatory efflux of Zn^2+^ into the MCV. This efflux is presumably carried out by ZntB, the main Zn^2+^ transporter in endosomes and phagosomes. Counterintuitively, *M. marinum* growth is also impaired in *zntB* KO cells, where MCVs accumulate less Zn^2+^. We also demonstrate that *M. marinum* senses toxic levels of Zn^2+^ and responds by upregulating its Zn^2+^ exporter CtpC, which supports bacteria survival under these restrictive conditions. Attenuation of *M. marinum* intracellular proliferation in *zntA* and *zntB* KO cells is accentuated in the absence of CtpC, confirming that mycobacteria face noxious levels of Zn^2+^. Altogether, we show for the first time that *M. marinum* infection induces a deleterious Zn^2+^ elevation in *D. discoideum*, which is counteracted by the bacteria with the induction of its Zn^2+^ exporter CtpC.

## INTRODUCTION

*Mycobacterium marinum* is a pathogenic bacterium that causes a tuberculosis-like infection in poikilotherms, such as fish and amphibians, in which it induces granulomatous lesions very similar to those caused in humans by its close relative *M. tuberculosis* (Mtb). *M. marinum* also opportunistically infects humans but, contrary to what happens during tuberculosis, infections by *M. marinum* are restricted to the skin and extremities due to their lower temperature (Aubry et al., 2017). At the cellular level of the host macrophage, the course of infection by both *M. marinum* and Mtb is strikingly similar, making *M. marinum* a experimentally versatile model to study pathogenic mechanisms of tuberculosis (Tobin and Ramakrishnan, 2008). Apart from fish and mammalian phagocytes, another well-established model host for *M. marinum* is the soil amoeba *Dictyostelium discoideum* (Cardenal-Munoz et al., 2017b), whose endocytic and cell-autonomous defence pathways are well conserved with humans (Dunn et al., 2017).

Upon phagocytosis by the host cell, *M. marinum* resides inside a phagosome that is rapidly and actively modified by the bacterium to divert from the canonical phagosome maturation pathway. It then becomes a replicative niche, the so-called *Mycobacterium*-containing vacuole (MCV). Like Mtb, *M. marinum* was thought to be a vacuolar pathogen. However, cumulating evidence indicates that both bacteria can also reside and proliferate in the cytosol of their host cells, where access to nutrients is not limited but bacteria are exposed to host cytosolic defences (Cardenal-Munoz et al., 2017b; Russell, 2001; Stamm et al., 2003). *M. marinum* starts damaging the MCV in the first hours of infection, before full rupture of the compartment releases the bacteria into the host cytosol at around 24-36 hours. Perforation of the MCV is achieved by secretion of the membrane-damaging peptide ESAT-6 through the mycobacterial ESX-1 Type VII Secretion System (T7SS) (Cardenal-Munoz et al., 2017a; Hagedorn et al., 2009; Lopez-Jimenez et al., 2018; Smith et al., 2008). The role of ESX-1-mediated secretion in MCV-to-cytosol translocation has also been shown for Mtb (Houben et al., 2012; Mittal et al., 2018; Simeone et al., 2012; van der Wel et al., 2007).

Infected cells use a vast repertoire of strategies to fight intracellular mycobacteria. For instance, *M. marinum* infection has been shown to induce iNOS (inducible nitric oxide synthase) gene expression in fish macrophages (Grayfer et al., 2011), and neutrophils kill these bacilli through NADPH-dependent mechanisms (Yang et al., 2012). In addition, *M. marinum* is targeted for digestion by the catabolic autophagy machinery of murine macrophages (Lerena and Colombo, 2011) and *D. discoideum* (Cardenal-Munoz et al., 2017a; Hagedorn et al., 2009). However, these mycobacteria have also evolved mechanisms of counter-defence against their hosts. They downregulate the iNOS levels (Thi et al., 2013) and suppress the production of reactive nitrogen intermediates (RNI), as well as they decrease the expression of NADPH oxidase components and reduce the production of reactive oxygen species (ROS) (Grayfer et al., 2011). Moreover, *M. marinum* avoids phagosome maturation by (i) modulating the composition of its cell wall (Robinson et al., 2007) or the MCV content in phosphatidylinositol-3-phosphate (PtdIns3P; (Koliwer-Brandl et al., 2019)), (ii) impairing the recruitment of the endosomal sorting complex required for transport (ESCRT)-0 component Hrs (Vieira et al., 2004) and (iii) avoiding accumulation of lysosomal enzymes (Cardenal-Munoz et al., 2017a; Hagedorn and Soldati, 2007; Koliwer-Brandl et al., 2019; Lerena and Colombo, 2011; Tan et al., 2006). It also blocks the autophagic flux, which would eventually deliver the bacteria into autolysosomes for killing (Cardenal-Munoz et al., 2017a; Lerena and Colombo, 2011).

Apart from the defence mechanisms mentioned above, little is known about other aspects of the chemical warfare between *M. marinum* and its hosts, and especially about the manipulation of transition metals by both parties. In the case of Mtb, it has been shown that immune cells deprive the bacteria from essential nutrients such as iron and manganese, whilst they intoxicate the mycobacteria by accumulating copper and zinc inside the MCV (Neyrolles et al., 2015; Pyle et al., 2017; Wagner et al., 2005; Wolschendorf et al., 2011). However, Mtb resists this metal fight with an arsenal of metal-binding proteins, oxidases and efflux transporters. For instance, Mtb captures Fe3^+^ from macrophages through its siderophore mycobactin (Sritharan, 2016), but it keeps low intracellular Cu^2+^ levels with its CtpV and MctB transporters and the Cu^2+^-binding metallothionein MymT (Gold et al., 2008; Ward et al., 2010; Wolschendorf et al., 2011). Mtb also resists the toxic free Zn^2+^ burst induced in human macrophages by exporting Zn^2+^ through the P-type ATPase CtpC (Botella et al., 2011), an efflux pump also present in *M. marinum* (Botella et al., 2012). On the host side, it has been shown that the mRNA levels of various Zn^2+^ transporters (i.e. ZIP8, ZnT1, ZIP1 and ZIP10) are altered in fish granulomas when compared to resting macrophages (Cronan et al., 2016). However, little is known about the conservation of these host versus pathogen strategies, and especially how mycobacterial CtpC and host Zn^2+^ transporters impact *M. marinum* infection in *D. discoideum*.

*D. discoideum* possesses four Zn^2+^ efflux transporters: ZntA, ZntB, ZntC and ZntD. ZntA locates to the contractile vacuole (CV), an organelle that regulates the osmotic and metal balance in the cell, and ZntB to organelles of the endosomal pathway (Barisch et al., 2018; Peracino et al., 2013). Whilst ZntA does not have any close homolog, ZntB shares homology with human ZNT1 and ZNT10, the latter being also present at early endosomes (Dunn et al., 2017). Similarly to their human homologs ZNT6 and ZNT7, located in the early secretory pathway (Dunn et al., 2017), *D. discoideum* ZntC and ZntD localize to the Golgi complex or recycling endosomes (Barisch et al., 2018). We wondered whether these Znts were involved in the resistance of *D. discoideum* to *M. marinum* infection. We demonstrate here that *D. discoideum* intoxicates *M. marinum* by inducing the expression and recruitment of ZntA and ZntB, but not ZntC nor ZntD, to the MCV. However, *M. marinum* resists the *D. discoideum* Zn^2+^ immune response by specifically expressing its Zn^2+^ exporter CtpC.

## RESULTS

### Free Zn^2+^ accumulates within intact *M. marinum*- containing vacuoles

It has been recently shown that, in *D. discoideum*, free Zn^2+^ accumulates inside (i) the CV network, (ii) zincosomes, endosomal compartments with lysosomal and post-lysosomal characteristics, and (iii) phagosomes containing beads and nonpathogenic bacteria such as *Escherichia coli* or *M. smegmatis* (Barisch et al., 2018; Buracco et al., 2017) (Fig. 1A). To test whether vacuoles containing the pathogenic bacterium *M marinum* also accumulated Zn^2+^, we monitored infected cells expressing the endosomal marker AmtA-mCherry and labelled with the pH-independent Zn^2+^ probe NBD-TPEA (Fig. 1B). Intact MCVs accumulated Zn^2+^ at all the stages of infection tested. On the contrary, visibly broken compartments appeared devoid of Zn^2+^, suggesting leakage due to mycobacteria-induced perforation of the MCV membranes. In fact, quantification of NBD-TPEA- labelled MCVs confirmed that, whilst wild-type (wt) MCVs were less positive for Zn^2+^ with progression of infection, intact MCVs containing *M. marinum* ∆RD1, a mutant lacking the ESX-1 secretion system and thus strongly attenuated in its capacity to induce membrane damage (Hagedorn et al., 2009; Lopez-Jimenez et al., 2018), remained positive for Zn^2+^ during the whole infection cycle (Fig. 1C).

**Fig. 1.**
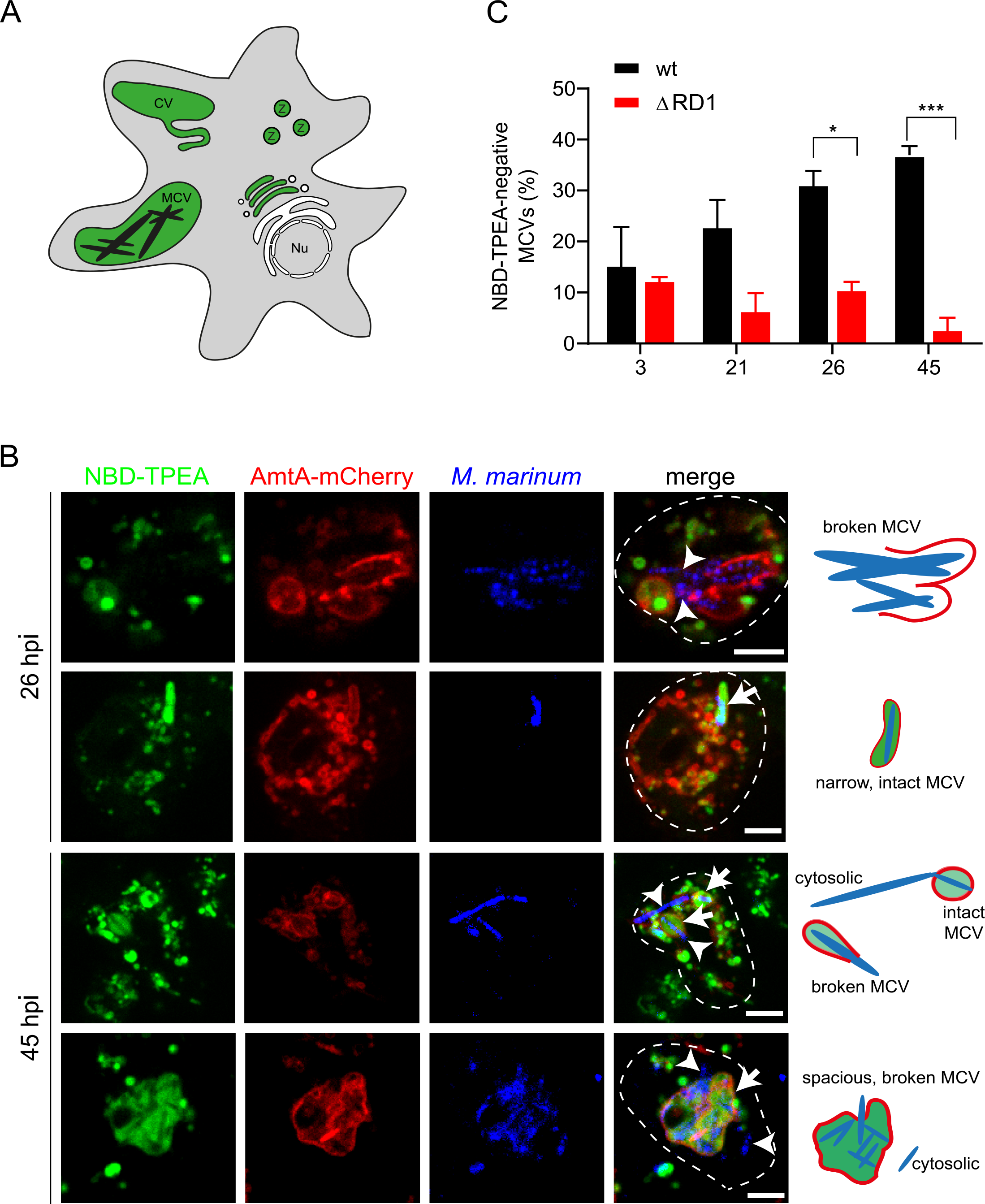
Free Zn^2+^ accumulates inside the MCV. **A.** Scheme depicting the different localizations of free Zn^2+^ in *D. discoideum*: zincosomes (Z), contractile vacuole (CV) and phagosomes (e.g. MCVs; (Barisch et al., 2018)). Nu, nucleus. **B.** Live imaging and illustrations of NBD-TPEA-treated cells expressing AmtA-mCherry. *M. marinum* was stained with Vibrant DyeCycle Ruby (blue) before imaging. Arrows point at Zn^2+^-positive MCVs; arrowheads label bacteria exposed to the *D. discoideum* cytosol. Scale bars, 5 μm. C. Percentage of NBD-TPEA-negative MCVs at different times post-infection with *M. marinum* wt and ∆RD1. Bars represent the mean and SEM of three independent experiments. About 300 wt and 200 ∆RD1 MCVs were counted in total. Statistical differences were calculated with a Tukey post hoc test after two-way ANOVA (\**p* < 0.05; \*\*\**p* < 0.001).

### *M. marinum* senses and reacts to toxic levels of Zn^2+^ by inducing its CtpC Zn^2+^ efflux pump

During infection of macrophages, Mtb is exposed to a burst of transition metals that induce in the bacteria the expression of efflux P-type ATPases such as CtpC, CtpG and CtpV (Botella et al., 2011; Tailleux et al., 2008). In particular, it has been shown that CtpC expression increases in Mtb during infection, which contributes to the bacteria detoxification from the accumulation of Zn^2+^ directed by the host (Botella et al., 2011). We wondered whether this was also the case in *M. marinum*. Increasing concentrations of Zn^2+^ lead to the corresponding increase in *ctpC* transcription (Fig. 2A). This upregulation by Zn^2+^ was CtpC-specific, since the mRNA levels of ZntA, another putative Zn^2+^-exporting P-type ATPase present in *M. marinum* but not in Mtb, did not change upon bacteria exposure to Zn^2+^ (Fig. 2A). Even the deletion of *ctpC* did not compensatorily induce the expression of ZntA, suggesting that, at least *in vitro*, CtpC is the major active *M. marinum* Zn^2+^ efflux transporter.

**Fig. 2.**
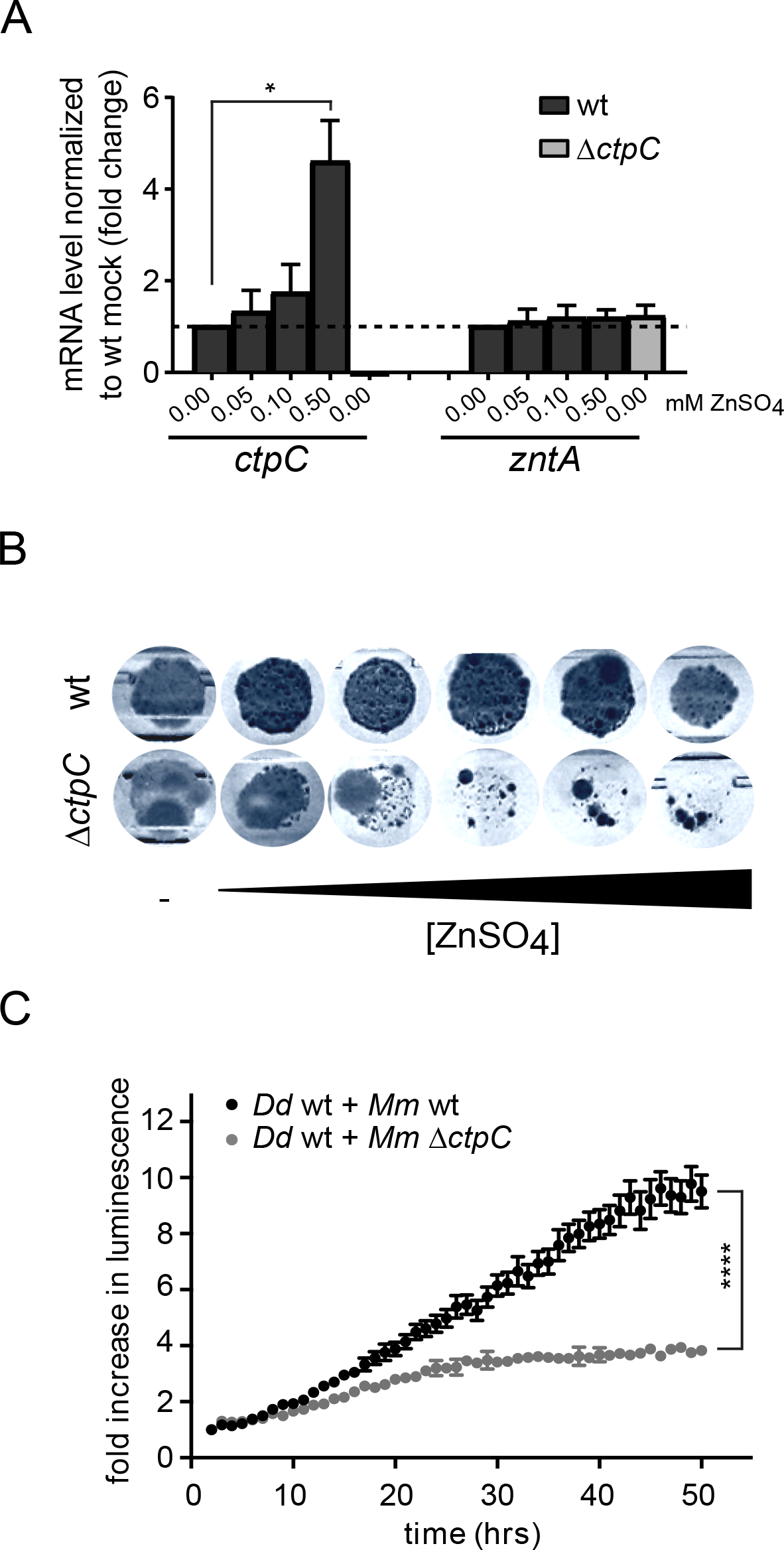
*M. marinum* senses and reacts to toxic levels of Zn^2+^ *in vitro* and during infection. **A.** Normalized mRNA levels of *ctpC* and *zntA* in GFP-expressing *M. marinum* grown in 7H9 with increasing concentrations of ZnSO_4_compared to bacteria grown in 7H9 without extra ZnSO_4_(depicted as dashed line). To confirm the successful deletion of *ctpC*, the mRNA levels in *M. marinum* ∆*ctpC* were tested as well. Shown are mean and standard deviations from two independent experiments. Statistical differences were calculated with an unpaired *t*-test (\**p* < 0.05). **B.** *M. marinum* wt and ∆*ctpC* were plated on 7H11 or 7H11 supplemented with increasing concentrations of ZnSO_4_ (0.125, 0.25, 0.5, 1 and 2.5 mM) as described in Material and Methods. Shown are the bacterial colonies obtained in one representative of three independent biological replicates, after seven days of incubation at 32°C. **C.** Cells were infected with *lux*-expressing *M. marinum* wt or ∆*ctpC*. The intracellular bacteria growth (in relative luminescence units [RLUs]) was monitored every hour. Shown is the fold increase in bacteria luminescence over time of one representative of three independent biological replicates. Error bars indicate the SEM of two technical replicates. Statistical differences were calculated with a Tukey test after two-way ANOVA (\*\*\*\**p* < 0.0001).

In addition, the growth of *M. marinum* ∆*ctpC* was impaired at high concentrations of Zn^2+^ (Fig. 2B). This was presumably due to the retention of toxic levels of Zn^2+^ within the bacteria, because high concentrations of other metals such as Mn^2+^ and Cu^2+^ did not differentially impact the growth of the ∆*ctpC* mutant (Fig. S1). Interestingly, the intracellular growth of *M. marinum* ∆*ctpC* was also notably impaired during infection of *D. discoideum* (Fig. 2C). We therefore conclude that, during infection of *D. discoideum*, *M. marinum* is exposed to toxic concentrations of Zn^2+^, and that the bacteria counteract this intoxication by inducing specifically its Zn^2+^ efflux pump CtpC.

### The *D. discoideum* Zn^2+^ transporters ZntA and ZntB localize to the *M. marinum* MCV

Since Zn^2+^ accumulated inside the *M. marinum* MCV (Fig. 1), we wondered whether the four already described *D. discoideum* Zn^2+^ efflux transporters ZntA, ZntB, ZntC and ZntD (Barisch et al., 2018), localized to the membranes of the compartment containing *M. marinum* and, therefore, were responsible for the Zn^2+^ enrichment within the MCV. Only ZntA and ZntB (Fig. 3A, B), but not ZntC nor ZntD (Fig. 3C, D), localized to the MCV, although with different kinetics. While ZntB was present throughout the infection cycle (Fig. 3B and E), ZntA was recruited to the MCV only at early time points (Fig. 3A and 3E). This suggests that ZntB is the main MCV Zn^2+^ transporter, but that ZntA also contributes to the import of Zn^2+^ into the MCV.

**Fig. 3.**
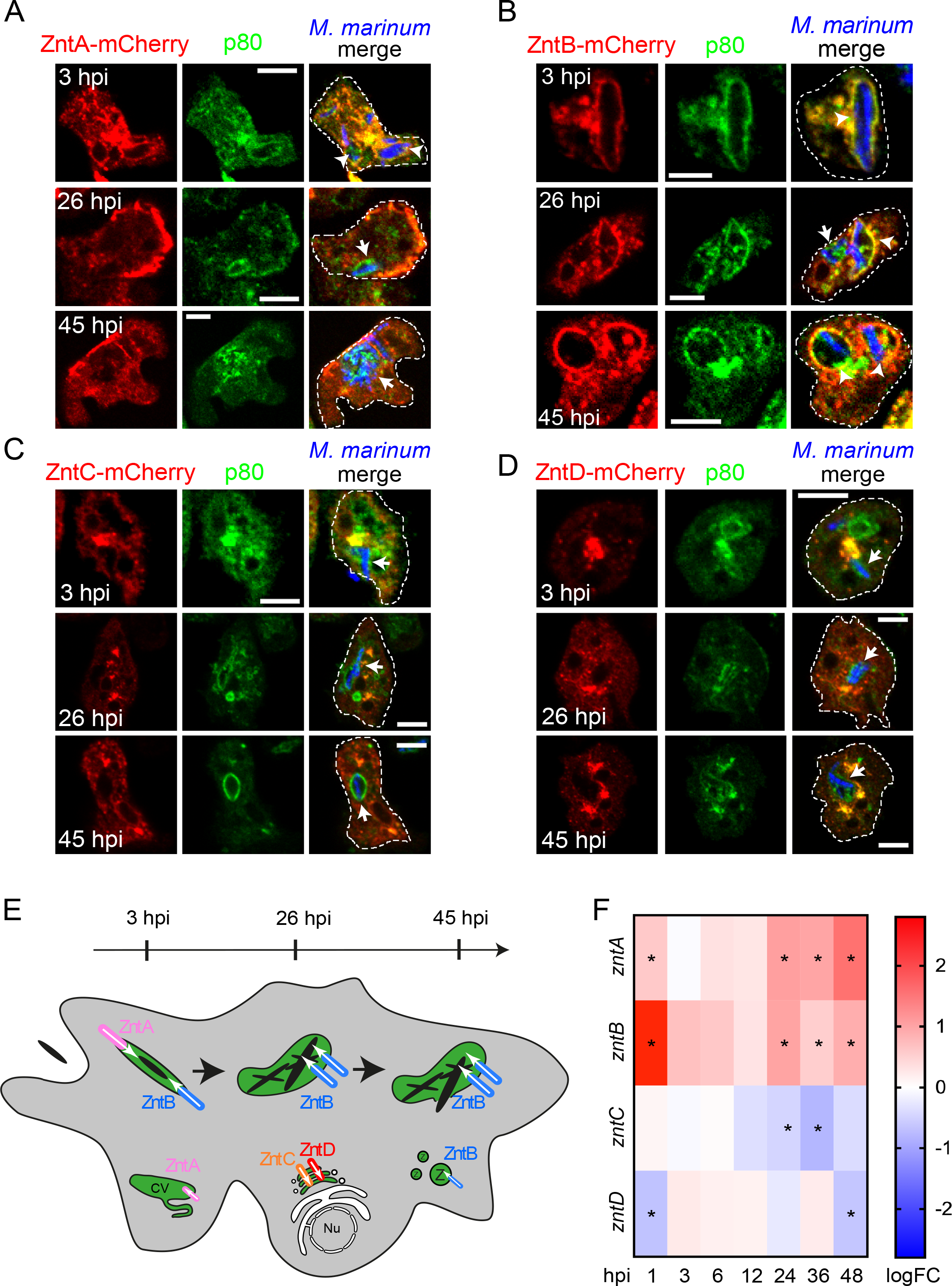
ZntA and ZntB transporters are recruited to the MCV and induced upon infection. **A-D.** Immunofluorescence staining at different times post-infection of ZntA-, ZntB-, ZntC- or ZntD-mCherry-expressing cells infected with GFP-expressing *M. marinum* (shown in blue). MCVs were visualized by staining for p80 (green). Arrowheads mark Znt-positive MCVs; arrows point to Znt-negative MCVs. Scale bars, 5 μm. **E.** Scheme depicting the localization of the four *D. discoideum* Znts during infection with *M. marinum*. **F.** Heat map representing the transcriptional data shown in Table S1. Cells were either infected with GFP-expressing *M. marinum* wt or mock-infected, and samples were collected at different hpi. The time points with statistically significant differential expression (p < 0.05) are marked with an asterisk. Colors indicate the strength of expression of *zntA-D* (in logFC) in infected cells compared to mock-infected: from dark red (highest expression) to dark blue (lowest expression).

### The expression of *Dictyostelium* Znts is altered during infection with *M. marinum*

As mentioned above, it has been described that *M. marinum* infection modulates the expression of various Zn^2+^ transporters in fish granulomas (Cronan et al., 2016). Since we observed a distinct pattern of recruitment of the *D. discoideum* Znts during infection, we wondered whether this pattern correlated with a differential regulation of the transporters at the transcriptional level. We analyzed recent RNA sequencing (RNA-Seq) data (A time-resolved transcriptomic analysis of the infection of *Dictyostelium* by *Mycobacterium marinum* reveals an integrated host response to damage and stress. N. Hanna, F. Burdet, A. Melotti, H. Hilbi, P. Cosson, M. Pagni, T. Soldati. To be submitted to bioRxiv), from which we specifically extracted the information related to the regulation of *D. discoideum zntA-D* during *M. marinum* infection (Fig. 3F and Table S1). Very interestingly, the expression of *zntA* and *zntB* was upregulated at very early, mid, and late times post-infection, with the highest expression levels found for *zntB* at 1 hour post-infection (hpi). On the contrary, the expression of *zntC* and *zntD* was not or only slightly affected during *M. marinum* infection. Altogether, we conclude that during *M. marinum* infection ZntA and ZntB are present at the MCV and their expression increases, presumably contributing to the accumulation of Zn^2+^ inside the MCV, whilst ZntC and ZntD are absent from MCVs and either not relevant for or even repressed during infection.

### Knockout of ZntA and ZntB impact on the concentration of Zn^2+^ inside the MCV

In order to define the relevance of ZntA and ZntB in the intoxication of *M. marinum* by Zn^2+^, we infected cells lacking one or the other transporter and monitored the levels of Zn^2+^ inside MCVs (Fig. 4A-D). Interestingly, compared to wt cells, the levels of Zn^2+^ in MCVs were lower in *zntB* knockout (KO) cells especially early during infection (3 hpi) (Fig. 4B, D) but higher in the *zntA* KO at later infection stages (46 hpi; Fig. A, C). This suggested that ZntB is the main exporter of Zn^2+^ into *M. marinum* MCVs, and that ZntB might compensate the absence of the contractile vacuole efflux transporter ZntA by augmenting the efflux of Zn^2+^ into the endosomal compartments and the MCVs (Fig 4E, F). These results are consistent with our previous findings showing the accumulation and the decrease of Zn^2+^ within phagosomes of *zntA* and *zntB* KO cells, respectively (Barisch et al., 2018). As shown for maturing phagosomes, Zn^2+^ might also be delivered to the MCV via fusion with zincosomes or other organelles positive for ZntC and ZntD.

**Fig. 4.**
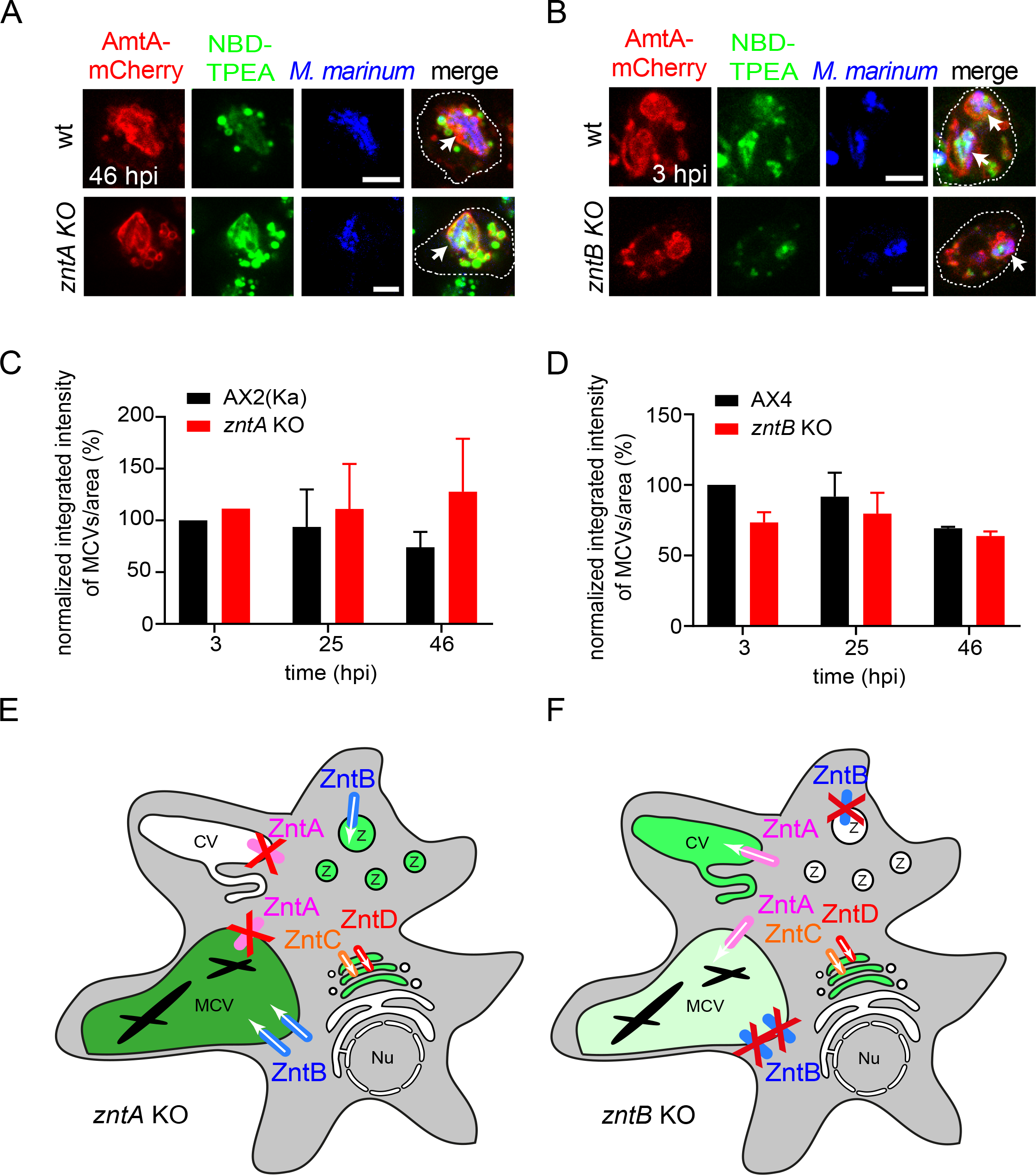
Zn^2+^accumulates in MCVs of cells lacking ZntA but decreases in *zntB* KO amoebae. **A-B.** Live imaging of NBD-TPEA-treated wt and *zntA* KO cells (**A**) or *zntB* KO cells (**B**) expressing AmtA-mCherry. *M. marinum* was stained with with Vibrant DyeCycle Ruby (blue) before imaging at 3 hpi (B) and 45 hpi, respectivley (A). Arrows label MCVs. Scale bars, 5 μm. **C-D.** Quantification of the normalized integrated intensity of NBD-TPEA inside MCVs per MCV area in AX2(KA) and *zntA* KO cells (**C**) or AX4 and *zntB* KO cells (**D**) infected with mCherry-expressing *M. marinum*. **E-F**. Schemes depicting the localization and concentration of Zn^2+^ in infected *zntA* (**E**) or *zntB* KO (**F**) cells. Z, zincosomes; CV, contractile vacuole; MCV, mycobacteria-containing vacuole; Nu, nucleus.

### Zn^2+^ restricts *M. marinum* during infection

We had previously shown that the accumulation of Zn^2+^ in phagosomes of *zntA* KO cells induced the more efficient killing of the non-pathogenic bacteria *E. coli* (Barisch et al., 2018). We wondered whether elevated Zn^2+^ would impact on *M. marinum* infection. Lack of ZntA lead to a decrease in the *M. marinum* intracellular load after 24 hpi (Fig. 5A), indicating that higher vacuolar Zn^2+^ was noxious to the bacteria (Fig. 4A and C). This was corroborated by the fact that the intracellular load of *M. marinum* ∆*ctpC*, which cannot expel Zn^2+^ from its cytosol, was even lower than that of *M. marinum* wt in *zntA* KO cells (Fig. 5A). This epistatic interaction between *M. marinum* ∆*ctpC* and *D. discoideum zntA* KO is a strong evidence of the role of Zn^2+^ during infection.

**Fig 5.**
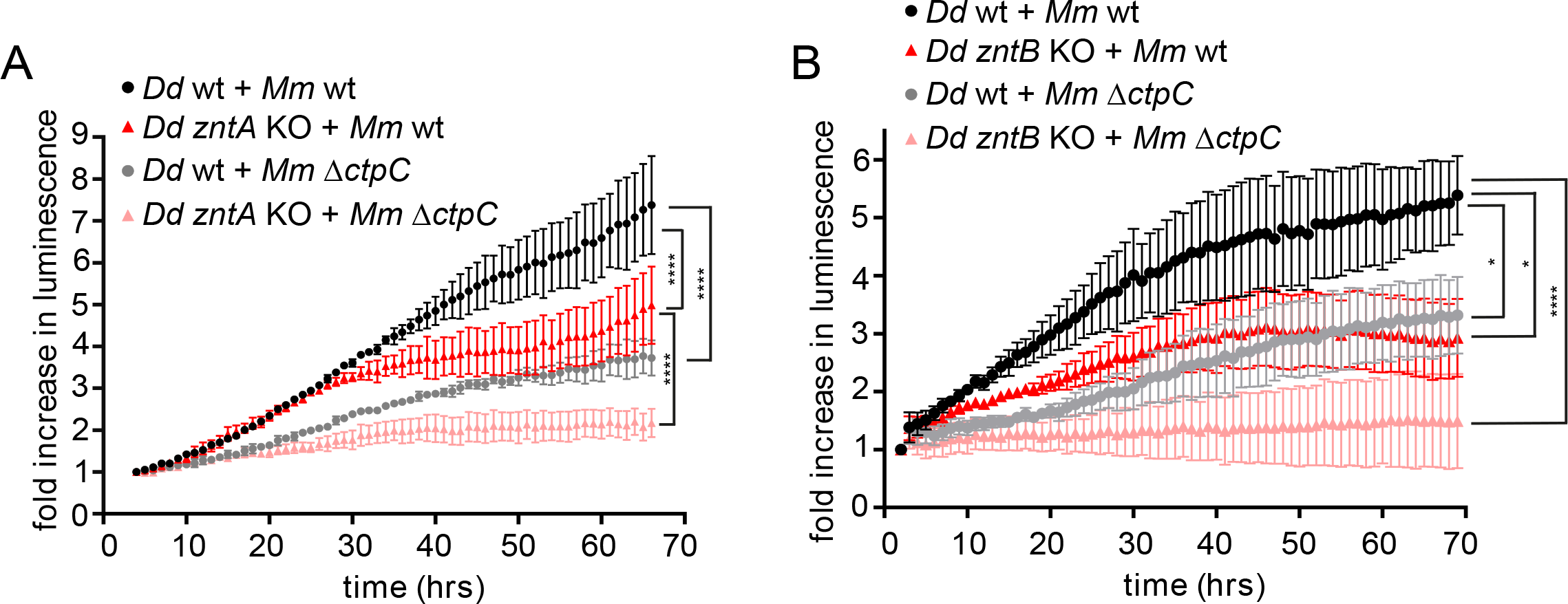
*M. marinum* intracellular growth is impaired in cells lacking ZntA or ZntB. **A.** AX2(Ka) and *zntA* KO cells were infected with *lux*- expressing *M. marinum* wt or Δ*ctpC*, and the intracellular bacteria growth (in RLUs) was monitored every hour. Shown is the fold increase in bacteria luminescence over time. Error bars indicate the SEM of three independent experiments. Statistical differences were calculated with a Tukey post hoc test after two-way ANOVA (\*\*\*\**p* < 0.0001). **B.** AX4 and *zntB* KO cells were infected with *lux*-expressing *M. marinum* wt or Δ*ctpC*, and the intracellular bacteria growth (in RLUs) was monitored every hour. Shown is the fold increase in bacteria luminescence over time. Error bars indicate the SEM of three independent experiments. Statistical differences were calculated with a Tukey post hoc test after two-way ANOVA (\**p* < 0.05; \*\*\*\**p* < 0.0001).

Enhanced killing of *M. marinum* by *zntA* mutant amoebae was also confirmed by the observed augmented growth of *zntA* KO cells on *M. marinum*- containing *Klebsiella pneumoniae* lawns (Fig. S2A, B). The formation of growth plaques by *D. discoideum* is a global indication of bacteria uptake, killing and digestion (Froquet et al., 2009). Therefore, since the *zntA* KO formed larger growth plaques on *M. marinum* than *D. discoideum* wt (Fig. S2A, B) and *M. marinum* intracellular load was lower in the *zntA* KO (Fig. 5A), we concluded that the lack of ZntA, and the consequent accumulation of Zn^2+^ within the MCV, leads to attenuation of *M. marinum* and therefore confers *D. discoideum* a growth advantage.

Intriguingly, although the concentration of Zn^2+^ inside *M. marinum* MCV was lower in cells lacking ZntB (Fig. 4B and D), the intracellular bacteria load also decreased with time in these cells (Fig. 5B). We noticed that whilst bacteria growth decreases inside *zntA* KO cells only after 24 hpi (Fig. 5A), the growth of *M. marinum* in *zntB* KO amoeabe was impaired from the beginning of the infection (Fig. 5B). Again, this epistatic interaction between *M. marinum* ∆*ctpC* and *D. discoideum zntB* KO is also a strong evidence of the role of Zn^2+^ during infection. At late stages of infection *M. marinum* escapes to the cytosol, and how the bacteria are restricted is imperfectly understood. We hypothesize that *M. marinum* growth inside *zntB* KO cells is restricted by other Zn^2+^-dependent killing mechanisms such as ROS production mediated by the activity of NADPH oxidases (DeCoursey et al., 2003; Hasegawa et al., 2000) or pathogen elimination by the autophagy pathway (Liuzzi et al., 2014). However, the *zntB* KO cells do not have a growth advantage on *M. marinum*- containing *K. pneumoniae* lawns (Fig. S2C, D) corroborating our previous results that food bacteria uptake, killing and digestion is unaffected in these cells (Barisch et al., 2018).

## DISCUSSION

Among other metals, immune cells use Zn^2+^ to fight intracellular microbes. Upon bacteria invasion, phagocytes manipulate their intracellular Zn^2+^ levels for nutritional immunity and/or metal intoxication purposes. In this manner, it has been reported that infected neutrophils weaken *Streptococcus pyogenes* growth by releasing the Zn^2+^ scavenger calprotectin (Makthal et al., 2017), while they enhance phagosomal Zn^2+^ levels for bacteria intoxication (Ong et al., 2014). Calprotectin also protects cells against other bacteria such as *Helicobacter pylori* (Gaddy et al., 2014), *Borrelia burgdorferi* (Lusitani et al., 2003) or *Staphylococcus aureus* (Corbin et al., 2008). As a different strategy, macrophages deliver Zn^2+^ to *E. coli*, *Salmonella enterica* serovar Typhimurium and Mtb-containing compartments to improve bacteria clearance (Botella et al., 2011; Kapetanovic et al., 2016). However, pathogens have evolved diverse mechanisms to counteract Zn^2+^-mediated defences. For instance, *Salmonella* evades Zn^2+^-containing vesicles by means of its pathogenicity island 1 (SPI-1) (Kapetanovic et al., 2016), while Mtb exports Zn^2+^ through its P-type ATPase CtpC (Botella et al., 2011). In addition, upon calprotectin-mediated chelation of Zn^2+^, *Salmonella* induces the expression of its Zn^2+^ importer ZnuABC, which ultimately supports bacterial growth (Liu et al., 2012). Here we focused on the Zn^2+^-mediated battle between *M. marinum* and its host *D. discoideum*.

*D. discoideum* has previously served as model to uncover the role of transition metals in the resistance to intracellular pathogens (Bozzaro et al., 2013; Buracco et al., 2017; Buracco et al., 2015; Peracino et al., 2006). For example, it has been shown that whilst Fe^2+^ participates in *D. discoideum* resistance to *M. avium* or *Legionella pneumophila* (Bozzaro et al., 2013; Buracco et al., 2017; Buracco et al., 2015; Peracino et al., 2006), Cu^2+^ or Zn^2+^ do not affect the infection by the latter (Buracco et al., 2017). Because little is known about the role of Zn^2+^ during *M. marinum* infection, (Cardenal-Munoz et al., 2017b) we used *D. discoideum*, a well-established host for *M. marinum* (Cardenal-Munoz et al., 2017b), to study Zn^2+^-mediated defence against these mycobacteria.

Contrary to the 24 transporters encoded in humans, *D. discoideum* only possesses eleven Zn^2+^ transporters: seven ZIP-like proteins (ZplA-G involved in the influx of Zn^2+^ from organelles or extracellular milieu to the cytosol) and four ZnTs (ZntA-D, responsible for the efflux of Zn^2+^ from the cytosol to organelles or extracellular milieu) (Dunn et al., 2017). ZntA is the main transporter in the CV and a key regulator of Zn^2+^ homeostasis in *D. discoideum*. When ZntA is not present, cells avoid toxic cytosolic concentrations of Zn^2+^ by reinforcing its pumping to the compartments of the endocytic pathway through the other transporters ZntB-D (Barisch et al., 2018). ZntB, is the main Zn^2+^ importer in lysosomes and recycling endosomes (Barisch et al., 2018). Interestingly, ZntA only located to early MCVs (Fig. 3A), whereas ZntB was present at the MCV throughout the intracellular cycle of *M. marinum* infection (Fig. 3B). Because cells lacking ZntB had decreased MCV Zn^2+^ levels, we hypothesized that ZntB is the main Zn^2+^ efflux pump during infection possibly directly contributing to the accumulation of Zn^2+^ inside the MCV (Fig. 4B-D). But in fact, in *zntB* KO cells, Zn^2+^ levels in the MCV were only moderately affected, leading to the conclusion that Zn^2+^ might be transferred into the MCV by the action of ZntA or by fusion with zincosomes as reported previously for the *M. smegmatis*-containing compartment (Barisch et al., 2018).

ZntC and ZntD were not detected at the MCV (Fig. 3C, D). In agreement with this, RNA-Seq data showed that the transcription of these two transporters was not altered, or even slightly reduced during *M. marinum* infection (Fig. 3F; Table S1; (Hanna et al., to be submitted to bioRxiv)). On the contrary, the mRNA levels of *zntA* and *zntB* increased upon infection with *M. marinum* (Fig. 3F; Table S1; (Hanna et al., to be submitted to bioRxiv)), consistently with their recruitment to the MCV (Fig. 3A, B). Strikingly, previous studies have shown that the expression of ZNT1, the homolog of ZntB, decreases in fish granulomas after two weeks of infection with *M. marinum*, but it increases in human macrophages after 18 and 72 hours of infection with Mtb (Botella et al., 2011; Cronan et al., 2016; Pyle et al., 2017). This suggests that distinct hosts respond differently to the infection and that, at least with regard to Zn^2+^ immunity, *D. discoideum* reacts to the invasion by pathogenic mycobacteria in a manner more similar to that of mammals than of fish.

We showed that *in vitro M. marinum* senses and reacts to elevated levels of Zn^2+^ by specifically inducing the expression of its Zn^2+^ efflux pump CtpC (Fig. 2A), which allows *M. marinum* wt, but not Δ*ctpC* to grow at high Zn^2+^ concentrations (Fig. 2B). Interestingly, *ctpC* is also induced during intracellular infection of *D. discoideum* (data not shown), substantiating that it experiences elevated Zn^2+^ concentrations. Furthermore, deletion of *ctpC* strongly attenuates *M. marinum* growth in *D. discoideum* (Fig. 2C). To our knowledge, this is the first study reporting the role of CtpC in *M. marinum* to resist Zn^2+^-mediated host defence.

In amoebae lacking ZntA or ZntB, the intracellular growth of *M. marinum* wt was lower than in wt hosts (Fig. 5A, B), and was even further reduced for Δ*ctpC*, leading to the conclusion that the mechanisms causing such a restriction are Zn^2+^-dependent but potentially different in the *zntA* KO and *zntB* KO mutants. Whilst KO of *zntA* drives the toxic accumulation of Zn^2+^ within MCVs (Fig. 4A, C), depletion of *zntB* leads to an attenuation of *M. marinum* that is probably mediated by other Zn^2+^-dependent killing mechanisms such as for example ROS generation or autophagy.

In summary, we conclude that *D. discoideum* restricts *M. marinum* using one or more Zn^2+^-dependent host defence mechanisms that are counteracted by induction of the mycobacterial Zn^2+^ exporter CtpC.

## MATERIALS AND METHODS

### *D. discoideum* strains, culture and plasmids

The *D. discoideum* material used in this study is listed in Table S2. *D. discoideum* (wt) strains [Ax2(Ka) and AX4] were grown at 22°C in HL5-C medium (Formedium) supplemented with 100 U/mL penicillin and 100 μg/mL streptomycin to prevent contamination. Overexpressors and KO cell lines were grown in medium with selective antibiotics hygromycin (25 µg/mL), G418 (5 µg/mL) and/or blasticidin (5 µg/mL)].

### *Mycobacterium* strains, culture and plasmids

The *M. marinum* strains used in the present study are listed in Table S3. *M. marinum* Δ*ctpc* was generated by specialized phage transduction as previously described (Table S3; (Koliwer-Brandl et al., 2019)), with modifications. The flanking regions of *ctpC* were amplified using the primer pairs oHK143/oHK144 and oHK145/oHK146 (Table S3). The PCR products were digested with *Alw*NI and *Van*91I, respectively, and ligated with *Van*91I-digested p0004S vector fragments. The resulting plasmid pHK42 and the DNA of the temperature-sensitive phage phAE159 were ligated after previous linearization with *Pac*I. For specialized transduction, high-titer phages were prepared in *M. smegmatis* mc^2^155 to yield to the *M. marinum* Δ*ctpC*::Hyg^r^ mutant. The phAE7.1 phage was used to remove the Hyg^r^ cassette (Koliwer-Brandl et al., 2019). Primers used for mutant verification are listed in Table S3 (oHK159, oHK160, oHK33 and oHK36).

Mycobacteria were cultured in Erlenmeyer flasks at 150 rpm at 32 °C in Middlebrook 7H9 (Difco) supplemented with 10% OADC, 0.2% glycerol and 0.05% Tween-80. Bacteria clumping was minimized by adding 5 mm glass beads into the flasks. Mutants and plasmid carriers were grown in medium with selective antibiotics [hygromycin (100 µg/mL), kanamycin (25 µg/mL) and/or ampicillin (100 µg/mL)].

### *M. marinum* growth on agar

*M. marinum* wt and Δ*ctpC* were grown in liquid 7H9-OADC- glycerol-tween at 32°C in shaking until an optical density at 600 nm (OD_600_) of approximately 0.5. Serial dilutions (dilution factor of 1:10 until 10^−4^) were performed and 5 µl of each dilution were plated on 7H11-OADC-glycerol-tween containing different concentrations (0, 0.125, 0.2, 0.5, 1 and 2.5 mM) of ZnSO_4_, MnCl_2_ and CuSO4. Pictures were taken after 7 days of incubation at 32°C.

### Infection of *D. discoideum*

Infections were performed as previously described (Hagedorn and Soldati, 2007). After washing off the extracellular bacteria, the infected cells were resuspended to a density of 1 x 10^6^ cells/mL in filtered HL5-C. 5 μg/mL of streptomycin and 5 U/mL of penicillin were added to prevent extracellular bacteria growth. The infected cells were then incubated at 25°C at 130 rpm, and samples were taken for analysis at the indicated time points.

#### Live imaging

To monitor the subcellular localization of Zn^2+^ during infection, infected cells were plated on μ-dishes (ibidi), medium was exchanged to Soerensen buffer and intracellular Zn^2+^ was stained with 5 µM NBD-TPEA (Sigma #N1040) for 30 min in the dark (Barisch et al., 2018). To stain unlabelled bacteria, Vibrant DyeCycle Ruby Stain (ThermoFisher) was used as previously described (Cardenal-Munoz et al., 2017a). Images were taken with an inverted 3i Marianas spinning disc confocal microscope using the 63× glycerol or 100× oil objectives.

To quantify the integrated intensity of NBD-TPEA inside MCVs, cells of wt (AX2(Ka) and AX4), *zntA* KO and *zntB* KO were infected with mCherry-expressing *M. marinum*. At the indicated time points, the infection was stained with NBD-TPEA and images were taken using an ImageXpress spinning disc confocal microscope (Molecular Devices). The integrated intensity per MCV area was assessed using an analysis pipeline created in MetaXpress (Molecular Devices).

#### Antibodies and immunofluorescence

The antibody against p80 (Ravanel et al., 2001) was purchased from the Geneva Antibody Facility (University of Geneva, Switzerland). The mCherry fluorescent signal was enhanced with rat mAb anti-RFP (Chromotek). As secondary antibodies, goat anti-rabbit, anti-mouse and anti-rat IgG coupled to Alexa488, Alexa546 (Thermo Fisher Scientific) or CF640R (Biotium) were used. Cells were fixed with cold MeOH as described previously (Hagedorn et al., 2006). Images were recorded with Zeiss LSM700 and LSM800 confocal microscopes and a 63×/1.4 NA or a 100×/1.4 NA oil-immersion objective.

#### RNA-Sequencing data

RNA-Seq data was gathered from Hanna *et al.* (Hanna et al., to be submitted to bioRxiv). Briefly, infection was carried out using *M. marinum*-expressing GFP as described above. Samples were taken over a time-course of infection and passed a fluorescence-activated cell scanning (FACS) sorter to obtain a homogeneous population of infected cells. The samples were sorted based on the emitted GFP fluorescence. Total RNA was extracted, rRNA from both infection partners was depleted, and rRNA-free samples were converted into cDNA libraries and sequenced. The resulting sequencing reads were mapped in parallel against the *D. discoideum* and *M. marinum* genome. Differential expression analysis of the time course was done by comparing infected cells versus Mock-treated using the R package limma.

#### qRT-PCR sample collection and analysis

To assess the effect of Zn^2+^ on *ctpC* and *zntA* expression, overnight *M. marinum* strains were exposed to various concentrations of ZnSO_4_ (0, 0.05, 0.1, and 0.5 mM) for two hours. Bacteria were harvested, RNA was extracted and cDNA was synthesized using BioRad iScript Kit. For each gene tested, the mean calculated threshold cycles (Ct) were averaged and normalized to the Ct of a gene with constant expression (*sigA*). The normalized Ct was used for calculating the fold change using the ΔΔC_t_ method. Briefly, relative levels of target mRNA, normalized with respect to an endogenous control (*sigA)*, were expressed as 2-ΔΔCt (fold), where ΔCt = Ct of the target gene - Ct of the control gene (*sigA*), and ΔΔCt = ΔCt of the studied set of conditions - ΔCt of the calibrator conditions, as previously described (Hanna et al., 2013).

#### Measurement of bacteria intracellular growth

Intracellular growth of *M. marinum* expressing bacterial luciferase was measured as previously described (Arafah et al., 2013). Briefly, three different dilutions of infected cells (between 0.5 and 2.0 × 10^5^ cells) were plated on non-treated, white F96 MicroWell™ plates (Nunc) covered with a gas permeable moisture barrier seal (Bioconcept). Luminescence was measured for around 70 h with 1 h intervals at a constant temperature of 25°C using a Synergy Mx Monochromator (Biotek).

#### Phagocytic plaque assay

The ability of *D. discoideum* AX2(Ka), *zntA* KO, AX4 and *zntB* KO to form plaques on *M. marinum* wt and Δ*ctpC* was assessed as described previously (Froquet et al., 2009). Briefly, 5 x 10^8^ mycobacteria were harvested and resuspended in 1.2 mL of 7H9-OADC-glycerol-tween containing a 1:10^5^ dilution of an overnight culture of *K. pneumoniae*. 50 μL of the suspension were added to the wells of a 24-well plate containing 2 mL of 7H11-OADC-glycerol-tween. Various dilutions of *D. discoideum* (i.e. 10, 100, 1000 and 10000 cells; for strains in AX4 background 3x more cells were used) were added to the bacterial lawn and plates were incubated at 25 °C for 7 days until plaques were visible. The logarithmic plaquing score was defined as follows: plaque formation in wells with 10 amoebae yielded a score of 1000; in the cases where cells did not grow at lower dilutions, they obtained the corresponding lower scores of 100, 10 and 1.

## ACKNOWLEDGEMENTS

We gratefully acknowledge the ACCESS imaging platform and the Bioimaging Center of the University of Geneva for their expert and friendly support. We are grateful to Dr Olivier Neyrolles for inspiring the project. The Soldati laboratory is supported by multiple grants from the Swiss National Science Foundation. Thierry Soldati is a member of iGE3 (www.ige3.unige.ch).

---

**Fig. S1.**
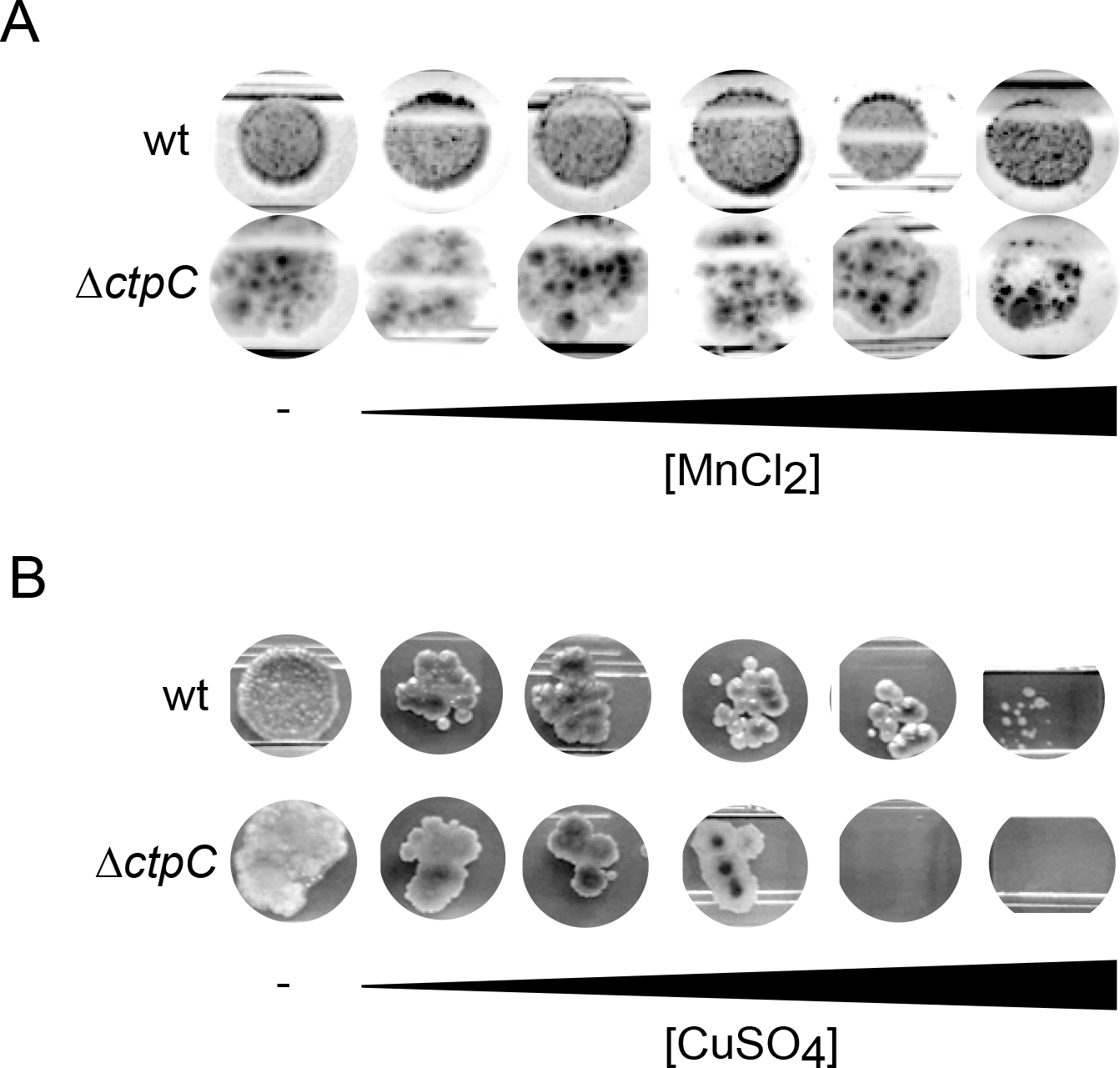
High concentrations of Mn^2+^ and Cu^2+^ do not differentially affect the growth of *M. marinum* ∆*ctpC*. **A-B.** GFP-expressing *M. marinum* wt and ∆*ctpC* were plated on 7H11 or 7H11 supplemented with increasing concentrations (0, 0.125, 0.25, 0.5, 1 and 2.5 mM) of MnCl_2_ (**A**) and CuSO_4_ (**B**) as described in Material and Methods. Shown are the bacterial colonies obtained in one representative of three independent biological replicates, after 7 days of incubation at 32°C.

**Fig. S2.**
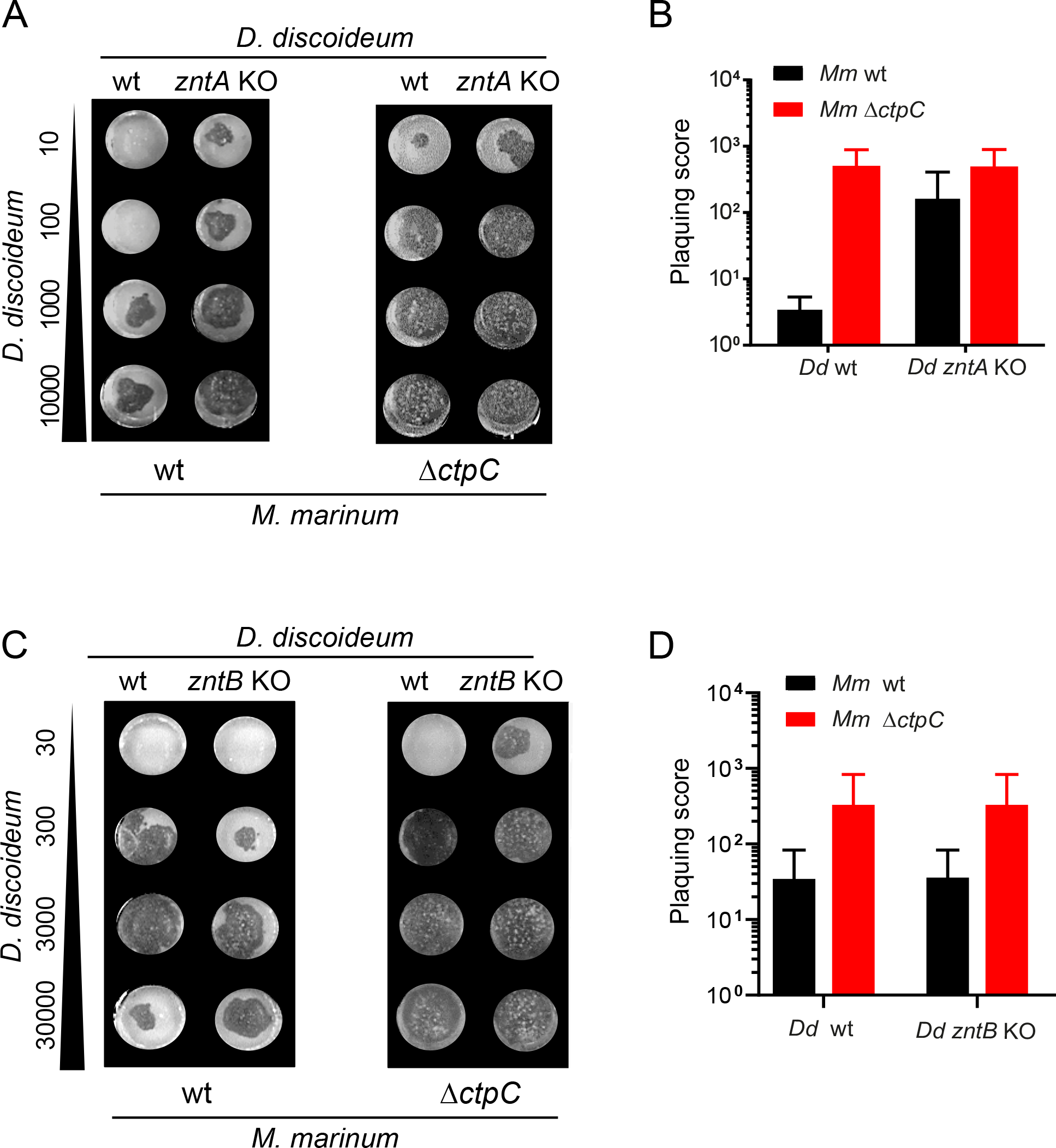
Growth of z*ntA* KO and *zntB* KO cells on *M. marinum/ K. pneumoniae* lawns. **A-D.** Plaque formation by AX2(Ka) wt and *zntA* KO (**A**) or AX4 wt and *zntB* KO cells (**C**) on GFP-expressing *M. marinum* wt and ∆*ctpC*, as described in Materials and Methods. Shown is one representative experiment from three independent biological replicates. **B** and **D**. Quantification of three independent experiments. Plaquing score was calculated as described in Materials and Methods

**Table S1.** Differential expression of *D. discoideum zntA-D* transporters during *M. marinum* infection (from (Hanna et al., to be submitted to bioRxiv)).

| Gene ID | Gene name | LogFC | Adjusted *p* value | Time point (hpi) |
| --- | --- | --- | --- | --- |
| DDB_G0283629 | *zntA* | 0.7376 | 4.86E-05 | 1 |
|  |  | -0.0593 | 0.9685 | 3 |
|  |  | 0.4146 | 0.4696 | 6 |
|  |  | 0.3508 | 0.3037 | 12 |
|  |  | 1.2701 | 3.70E-08 | 24 |
|  |  | 1.2005 | 1.55E-07 | 36 |
|  |  | 1.8154 | 1.41E-13 | 48 |
| DDB_G0282067 | *zntB* | 2.7400 | 2.73E-23 | 1 |
|  |  | 0.8484 | 0.1141 | 3 |
|  |  | 0.7464 | 0.2132 | 6 |
|  |  | 0.3817 | 0.4262 | 12 |
|  |  | 1.2188 | 0.0001 | 24 |
|  |  | 0.6640 | 0.0399 | 36 |
|  |  | 1.0877 | 0.0007 | 48 |
| DDB_G0269332 | *zntC* | 0.1333 | 0.5892 | 1 |
|  |  | -0.0942 | 0.9518 | 3 |
|  |  | 0.0540 | 0.9718 | 6 |
|  |  | -0.4570 | 0.2011 | 12 |
|  |  | -0.5898 | 0.0260 | 24 |
|  |  | -0.9956 | 0.0001 | 36 |
|  |  | -0.5010 | 0.0519 | 48 |
| DDB_G0291141 | *zntD* | -0.8446 | 0.0009 | 1 |
|  |  | 0.2956 | 0.8772 | 3 |
|  |  | 0.2080 | 0.8716 | 6 |
|  |  | 0.1851 | 0.7546 | 12 |
|  |  | -0.3203 | 0.3177 | 24 |
|  |  | 0.2661 | 0.4402 | 36 |
|  |  | -0.7934 | 0.0065 | 48 |

*D. discoideum* wt was either infected with GFP-expressing *M. marinum* wt or mock-infected. Samples were collected at different times post-infection (1, 3, 6, 12, 24, 36 and 48 hpi) and sorted for enrichment in infected cells (see Materials and Methods). Shown are the logarithmic fold-changes (LogFC) in gene expression between infected and mock-infected samples.

**Table S2.** *D. discoideum* material used for this study.

| **Strains** | **Relevant characteristics** | **Source/Reference** |
| --- | --- | --- |
| Ax2(Ka) | wt, parental strain of the *zntA* KO |  |
| Ax2(Ka) *zntA* KO | BSD<sup>r</sup> | (Barisch et al., 2018) |
| AX4 | wt, parental strain of the *zntB* KO |  |
| AX4 *zntB* KO | BSD<sup>r</sup> | REMI library, Prof. Christopher Thompson (Barisch et al., 2018) |
| **Plasmids** |  | **Source/Reference** |
| ZntA-mCherry | pDM1044-*zntA*, Hyg<sup>r</sup> | (Barisch et al., 2018) |
| ZntB-mCherry | pDM1044-*zntB*, Hyg<sup>r</sup> | (Barisch et al., 2018) |
| ZntC-mCherry | pDM1044-*zntC*, Hyg<sup>r</sup> | (Barisch et al., 2018) |
| ZntD-mCherry | pDM1044-*zntD*, Hyg<sup>r</sup> | (Barisch et al., 2018) |
| AmtA-mCherry | pDM1044-*amtA*, Hyg<sup>r</sup> | (Barisch et al., 2015) |

**Table S3.**
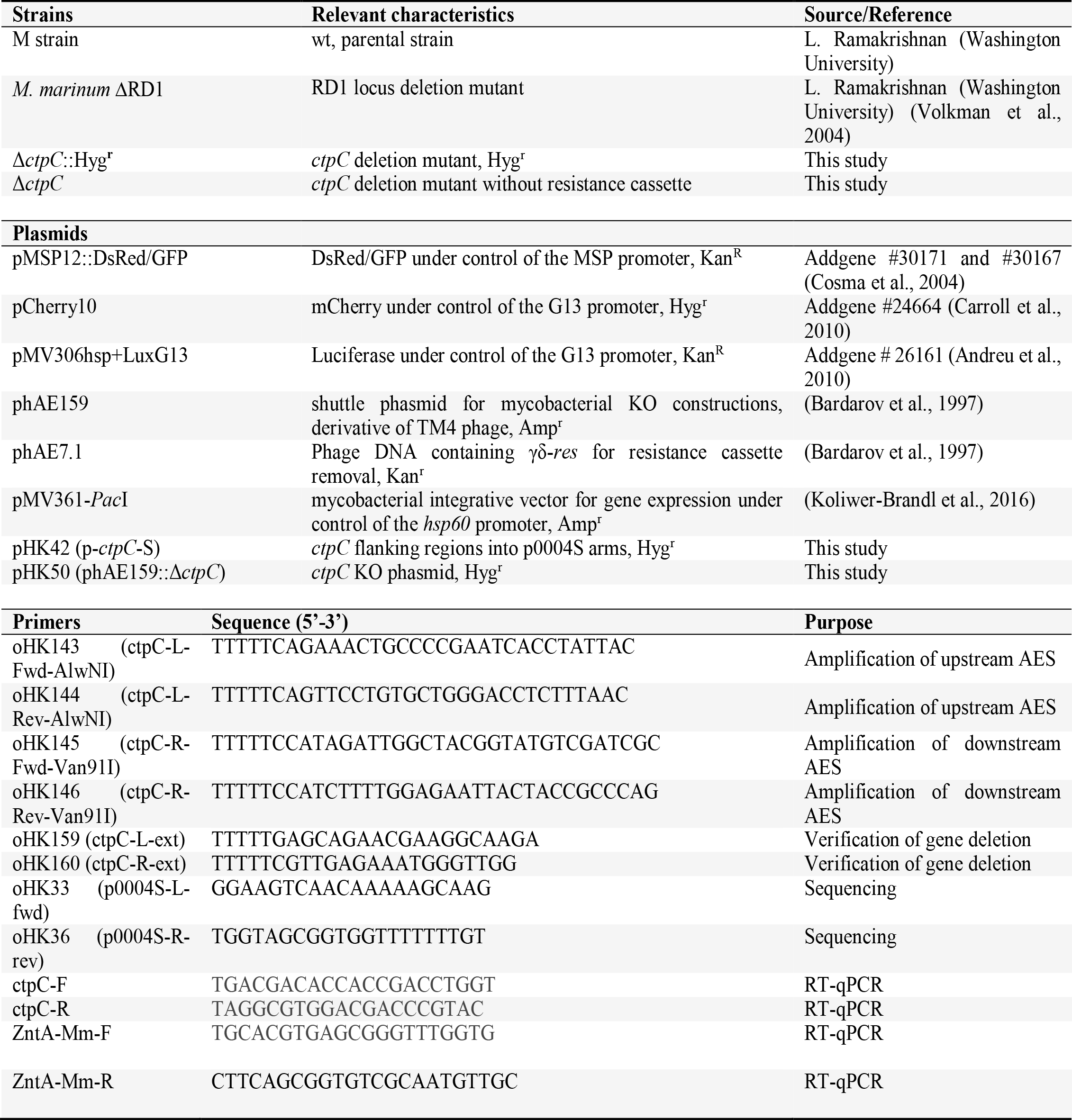
*M. marinum* material used in this study.

